# The Female Heart: Sex Differences in the Dynamics of ECG in Response to Stress

**DOI:** 10.1101/368845

**Authors:** Tricia Adjei, Jingwen Xue, Danilo P. Mandic

**Author notes:** Correspondence: Tricia Adjei.

## Abstract

Sex differences in the study of the human physiological response to mental stress are often erroneously ignored. To this end, we set out to show that our understanding of the stress response is fundamentally altered once sex differences are taken into account. This is achieved by comparing the heart rate variability (HRV) signals acquired during mental maths tests from ten females and ten males of similar maths ability; all females were in the follicular phase of their menstrual cycle. For rigour, the HRV signals from this pilot study were analysed using temporal, spectral and nonlinear signal processing techniques, which all revealed significant statistical differences between the sexes, with the stress-induced increases in the heart rates from the males being significantly larger than those from the females (p-value=2.2×10^−3^). In addition, mental stress produced an overall increase in the power of the low frequency component of HRV in the males, but caused an overall decrease in the females. The stress-induced changes in the power of the high frequency component were even more profound; it greatly decreased in the males, but increased in the females. We also show that mental stress was followed by the expected decrease in sample entropy, a nonlinear measure of signal regularity, computed from the males’ HRV signals, while overall, stress manifested in an increase in the sample entropy computed from the females’ HRV signals. This finding is significant, since mental stress is commonly understood to be manifested in the decreased entropy of HRV signals, however, the significant difference (p-value=2×10^−9^) in the changes in the entropies from the males and females highlights the pitfalls in ignoring sex in the formation of a physiological hypothesis. Furthermore, it has been argued that oestrogen attenuates the effect of catecholamine stress hormones; the findings from this investigation suggest for the first time that the conventionally cited cardiac changes, attributed to the fight-or-flight stress response, are not universally applicable to females. Instead, this pilot study provides an alternative interpretation of cardiac responses to stress in females, which indicates a closer alignment to the evolutionary tend-and-befriend response.

## 1 INTRODUCTION

The effects of stress on heart rate (HR) and heart rate variability (HRV) are considered to be well defined and long established. General physiological stress is understood to cause an increase in HR, consequently decreasing HRV (Houtveen *et al.*, 2002), whilst physical stress has been widely reported to cause an increase in the power of the low frequency (LF) component of HRV signals, and a decrease in the power of the high frequency component (Montano *et al.*, 1994). It is therefore commonly conjectured, though not without controversy, that the LF component of HRV (0.04-0.15 Hz) reflects the activity of the sympathetic nervous system (SNS), while the HF component (0.15-0.4 Hz) reflects the activity of the parasympathetic nervous system (PNS). The controversy surrounding this conjecture was highlighted in several studies where the effects of mental stress on the LF and HF components of HRV were found to be neither consistent, nor correlate with the trends seen in physical stress (Berntson *et al.*, 1994). However, despite the uncertainty over the physiological interpretations of the LF and HF components, and whether or not they respectively represent the activities of the SNS and PNS (Berntson *et al.* (1994), Eckberg (1997), Berntson and Cacioppo (2004), Billman (2013) and von Rosenberg *et al.* (2017)), it comes as a surprise that until recently, the theories regarding the relationships between stress, HR and HRV were developed without accounting for, or appreciating sex differences. To this end, this work aims to quantify and demystify the effects of sex differences on the dynamics of ECG, in a proof-of-concept study based on ECG recordings from participants who have been subjected to a mental stressor.

The commonly adopted theory of the stress response stems from a seminal book by Walter Cannon, in which it was hypothesised that, irrespective of sex, the evolutionary purpose of the human stress response is to prepare the body for fight or flight when faced with a threat (Cannon, 1915). It is generally accepted that reaching either the fight or flight stage requires the energisation of the body through stress hormones, for example, by increasing HR, arousal and the respiratory rate. However, long-term exposure to these hormones is known to lead to pathologies, such as heart disease, insomnia and hyperventilation (NHS Derbyshire Health Psychology Service, 2012). Chronic stress is also linked to depression; it is known that chronic stress causes structural degeneration in the pre-frontal cortex, which is a major risk factor for depression (Mah *et al.*, 2016). It would therefore be expected that the correlations between many stress induced pathologies, such as heart disease and depression, would be high; instead, the statistics indicate no such correlation (Åhs *et al.*, 2009).

Whilst females have the highest incidence of depression, the highest incidence of cardiac pathologies is seen in males. For example, in 2014, 4.3% of UK females had experienced depressive episodes, compared to 3.2% of males (NHS Digital, 2016). In contrast, in the year 2013/14, the number of UK males admitted to hospital with heart related diseases was 30.3% higher than the corresponding number of females (British Heart Foundation, 2015). Such stark statistical differences have led to further investigation into the causes of this disparity. An insightful overview of sex differences in the response to stress by Verma *et al.* (2011) reports many hormonal, neuroanatomical and cognitive differences between the sexes; it concludes that males and females exhibit distinct psychological and biological differences in their responses to stress. Similarly, Ramaekers *et al.* (1998) assessed sex differences in cardiovascular dynamics in response to stress, by analysing 24-hour-long HRV signals, recorded from 135 females and 141 males aged between 18 and 71 years. They reported that the absolute powers of the LF components of HRV from the males, regardless of age, were significantly larger than those from the females aged under 40, but were not significantly larger than those from the females aged over 40 (Ramaekers *et al.*, 1998). This age-dependence has indicated that the possible sex difference in cardiovascular dynamics is due to the effects of the menstrual cycle, and in particular, oestrogen (Ramaekers *et al.*, 1998). It is now hypothesised that oestrogen enhances parasympathetic control of the heart (Dart *et al.*, 2002), which in turn means that premenopausal females will experience enhanced parasympathetic control compared to males and postmenopausal females. This gives a wide scope for the study of the effect of oestrogen on cardiovascular dynamics, which promises to dramatically alter the understanding of the stress response, a subject of this work.

At present, the effect of hormones on the cardiovascular response to stress is considered to be related to the activation of the sympathetic nervous system (SNS), which mediates the release of catecholamines and glucocorticoids (Lundeberg, 2005). Many researchers have posited that once a stressor diminishes, the parasympathetic nervous system (PNS) takes dominance over the SNS (Figueroa-Fankhanel, 2014); the PNS is known to restore vital functions to their rest state (Thayer *et al.*, 2009). However, the theory that the SNS and PNS have an antagonistic or reciprocal relationship has also become controversial (Billman, 2013). An often cited review by Eckberg (1997) assesses SNS and PNS dynamics, and reports many parallel activations of the SNS and PNS, and that there is no definitive physiological evidence to suggest that the SNS and PNS must behave reciprocally. Eckberg (1997) describes the tendency to assign a reciprocal relationship to the SNS and PNS as philosophical, as opposed to physiological. Another study often cited to support nonreciprocal SNS and PNS dynamics is Berntson *et al.* (1994), who assessed the SNS and PNS responses in 10 participants subjected to mental stress, and reported slightly positive correlations between the activities of the SNS and PNS. They concluded that certain stressors may elicit autonomic responses which are specific to individuals, whilst others elicit common SNS and PNS responses. Yet, despite the popularity of the paper, it is often overlooked that all 10 of the subjects recruited by Berntson *et al.* (1994) were young females. In light of the work by Dart *et al.* (2002), a study cohort made up entirely of young females, without an account of their menstrual cycle phase, could explain why the results reported by Berntson *et al.* (1994) differed from similar studies. In summary, conflicting findings regarding the cardiovascular responses to mental stress point to a lack of clarity in the understanding of the stress response; this motivated us to ask whether this lack of clarity was simply due to conflating the male and female responses to stress?

To provide a conclusive answer to this question, we set out to assess the cardiovascular dynamics of young males and young females during a mental stress task; this was achieved in a rigorous, quantitative, and reproducible way, by applying state-of-the-art signal processing techniques to HRV signals. We make no attempt to correlate the LF and HF components of HRV to the SNS and PNS, but to identify differences in the temporal, spectral and nonlinear characteristics of the recorded signals. The changes in heart rate will be used to characterise signals temporally, whilst spectral characterisation is achieved through the computation of the powers of the LF and HF components of HRV, and nonlinear characterisation is accomplished using the nonparametric sample entropy (SE) method. Our results show conclusively that males and females in the follicular phase of their menstrual cycle exhibit significantly different cardiovascular responses to mental stress.

## 2 MATERIALS AND METHODS

### 2.1 Subjects

Our study cohort consisted of 10 males and 10 females, with respective mean ages of 28.6 (standard deviation: 5.6, range: 23 to 38) and 23.3 (standard deviation: 1.3, range: 21 to 26). The size of the dataset used is validated by a statistical study (Ristic-Djurovic *et al.*, 2018), which reported that nine is the minimum recommended number of subjects to draw statistically significant conclusions from a biomedical study.

The calendar method was used to confirm that all female subjects were in the follicular phase of their menstrual cycle; the follicular phase precedes ovulation, and is characterised by increasing oestrogen levels (Dart *et al.*, 2002).

The experimental procedure was explained to the subjects, both verbally and in writing, and all subjects gave their full consent to take part in the study. Ethics approval was granted by the Joint Research Office at Imperial College London (reference IRREC_12_1_1).

### 2.2 Experimental Procedure

The electrocardiogram (ECG) was recorded from each subject as they sat at rest for 15 minutes, and during a 15-minute mental maths test in pairs (subjects competed against sex-matched opponents with similar mathematical abilities and proficiency in the English language). There was a one-minute interval between the rest and test periods, and to induce further mental stress, the subjects were told that their performance in the maths test would be recorded.

### 2.3 Data Acquisition

The subjects’ ECGs were recorded by a custom-made data logger, called the iAmp (Kanna *et al.*, 2018), using adhesive surface electrodes placed in the Lead I ECG configuration, and at a sampling frequency of 1000 Hz.

All signal processing was performed in the Matlab programming environment. The ECG signals from the subjects were segmented into epochs, corresponding to the rest and mental maths test, before HRV was derived. The R-waves in the ECGs were extracted and interpolated at 4 Hz to derive HRV, using the robust algorithm introduced in Chanwimalueang *et al.* (2015). Temporal, spectral and nonlinear signal processing techniques were applied to the epochs of HRV to extract the four cardiac metrics described below.

#### 2.3.1 Temporal Analyses

The HR from the HRV signals was computed from a one-minute sliding window, with a one-second increment, where xi denotes an HRV data point in the windowed signal, and n designates the number of data points in the window, to yield

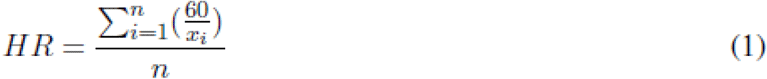

#### 2.3.2 Spectral Analyses

The powers of the LF and HF bands in HRV were computed as the powers of the 0.04-0.15 Hz and 0.15-0.4 Hz bands respectively, and were normalised by the power of the 0.04-0.5 Hz band, N_p_, to give

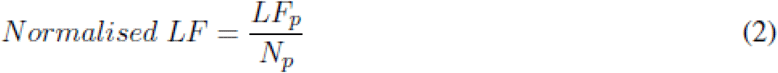

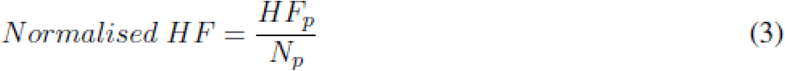

where the respective powers of the LF and HF bands are denoted by LF_p_ and HF_p_.

This new N_p_ power band of interest was used in Equations 2 and 3 for the normalisation, instead of the total power for the following reasons: (i) it has long been known that the frequencies below 0.04 Hz have no clear physiological interpretation (Malik *et al.*, 1996), (ii) heart beats are produced at a frequency of at least 1 Hz, which means the effective sampling frequency of HRV is also 1 Hz; the Nyquist theorem therefore suggests all useful information in HRV is contained below 0.5 Hz (Kuusela, 2004).

The mean normalised powers of the LF and HF bands were computed for both the rest and mental maths epochs, for each subject. The mean normalised powers of the LF and HF bands will be simply referred to as LF and HF, respectively.

#### 2.3.3 Nonlinear Analyses

Numerous studies have investigated the effect of stress on the structural complexity within HRV signals. Central to the analysis of signal complexity is the complexity loss theory as introduced by Goldberger *et al.* (2002), which posits that any perturbations within a physiological system, such as those caused by stress, constrain the system. Goldberger *et al.* (2002) hypothesise that the signals recorded from a constrained physiological system will exhibit reduced structural complexity. Signal complexity is interpreted through the regularity of a signal and is measured using entropy algorithms; equating structural complexity to signal regularity justifies the use of entropy to analyse signal complexity, as entropy algorithms are widely used measures of regularity. However, a truly complex system is neither completely regular nor irregular (Tononi *et al.*, 1998), which possibly renders entropy an inadequate measure of complexity (Goldberger *et al.*, 2002).

Nevertheless, many studies which use entropy to assess the effects of stress on signal complexity have concluded that stress reduces the complexity within cardiac signals, supporting the complexity loss theory. Vuksanovic and Gal (2007), Williamon *et al.* (2013) and Chanwimalueang *et al.* (2016) employed the sample entropy method to analyse the effect of mental stress on the structural complexity in HRV signals, and found that mental stress led to a decrease in entropy. Bornas *et al.* (2006) employed sample entropy to analyse ECG signals from flight phobics, and found that sitting in a flight simulator resulted in a reduction in the entropy of their ECG. Given the choice of the sample entropy method over other entropy measures in these relevant studies, we here apply sample entropy to assess sex differences in the effects of stress on the structural dynamics of HRV signals.

The sample entropy algorithm was introduced by Richman and Moorman (2000), and is the negative natural logarithm of the likelihood that two similar segments of data will remain similar, within a given tolerance, if the lengths of the segments are increased by one data point; see Step 1 to Step 7 in Algorithm 1 (Costa *et al.*, 2002, Costa *et al.*, 2005 and Song *et al.*, 2012).

The sample entropy analyses in this study were undertaken in a five-minute sliding window, with a one-second increment; mean SE values for every subject for the two experiment epochs were computed.

#### 2.3.4 Statistical Analyses

The percentage changes in the above described cardiac metrics, from the rest to maths epochs, were used to compare the male and female cardiovascular reactions to mental stress. The use of percentage changes enables inter-subject statistical comparisons, whilst the choice of the statistical test employed was dependent on the distribution of percentage changes. For example, Student’s t-test was used to compare the results which followed a normal distribution, and the Wilcoxon rank-sum was used to compare the results which did not follow a normal distribution. A significance level of 0.01 was assumed in all tests.

The analysis was performed within a five-minute sliding window, with a one-second increment. This window duration is the minimum recommended length to fully capture cardiac dynamics (Malik *et al.*, 1996).

## 3 RESULTS

Figure 1 A shows that mental stress induced a significantly greater increase in HR in the males, compared to the females (Wilcoxon rank-sum: p-value=2.2 x10^−3^). The median increase in HR in the males was 11%, with a range of +10% to +22%; in contrast, the median percentage change in the females was an increase of only 5%. The changes in HR in the females were also less consistent, with a range of −7% to +13%. The median changes in the cardiac metrics and the p-values indicating the significance of the differences between the male and female responses are shown in Table 1.

**Table 1.**
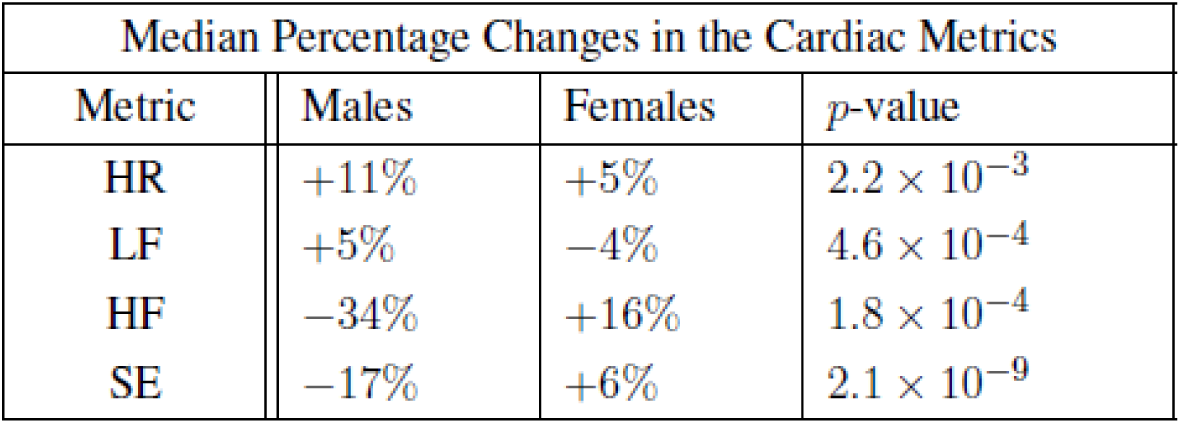
The percentage changes in the cardiac metrics from the rest epochs to the maths epochs

**Figure 1.**
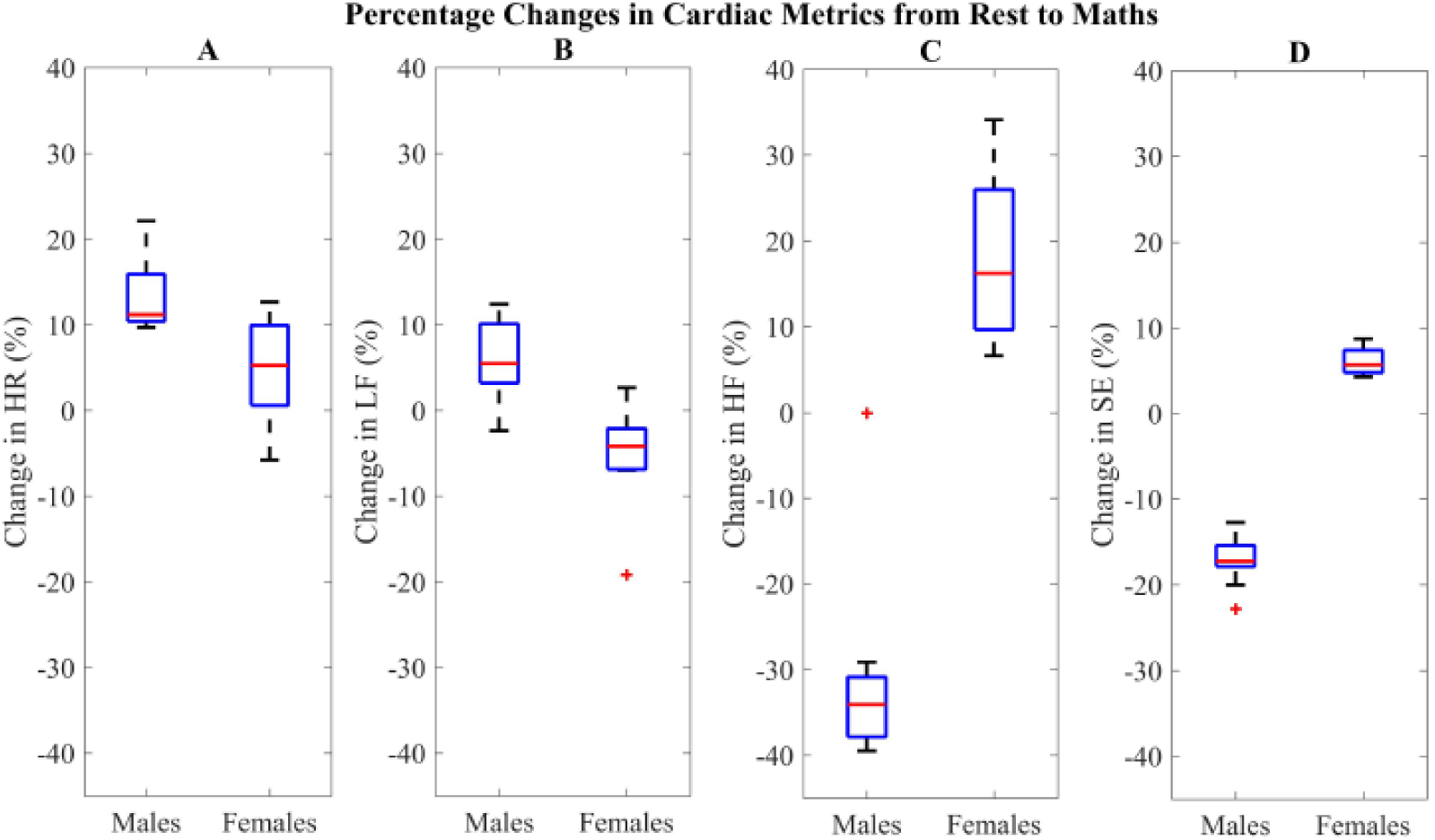
Temporal, spectral and non linear analyses of HRV. A) Distribution of percentage changes in HR from the rest to maths epoch. B)Distribution of percentage changes in HF from the rest to maths epochs. D)Distribution of percentage changes in SE from the rest to maths epochs.

The results from the spectral analyses show a non-reciprocal relationship between LF and HF; the median changes in LF and HF in the males were a respective 5% increase and a 34% decrease, whilst the corresponding changes in the females were a 4% decrease in LF and a 16% increase in HF.

Figures 1 B and 1 C illustrate the sex differences in the cardiac responses to mental stress. The percentage changes in LF from the males were significantly different to those from the females (t-test: p-value=4.6 x10^−4^), and were more varied, with a range of −2% to +12%; those from the females ranged from -7% to +3%. In addition, the percentage changes in HF were far more distinguishing (Wilcoxon rank-sum: p-value=1.8 x10^−4^). The changes in HF in the males were more concentrated, with a range of −40% to −30%, whereas the changes in the females ranged from +7% to +34%. Figure 1 D shows the findings from the nonlinear analysis, which revealed the most significant difference between the male and female cardiac responses to mental stress (t-test: p-value=2.1 x10^−9^). The median percentage change in SE in the males was a 17% decrease, with a range of −20% to −13%. The median percentage change in the females was a 6% increase in SE, with a range of a +4% to +9%.

## 4 DISCUSSION

The changes in HR shown in Fig. 1 A provide the first conclusive evidence that the expected increase in HR in response to stress is not universal; while every male experienced an increase in HR of at least 10%, only half of the females experienced increases of at least 5%. The results from this study therefore confirm that the effect of stress on cardiac dynamics can differ substantially between males and females; this calls for a re-evaluation of our understanding of how stressful events affect cardiac dynamics in females in the follicular phase of their menstrual cycle. A similar sex difference has previously been reported in Tousignant-Laflamme *et al.* (2005) in relation to the effects of pain (a form of physiological stress) on HR. It was found that whilst the correlation between pain intensity and HR was positive in males, no such correlation existed in the females (Tousignant-Laflamme *et al.*, 2005). However, Tousignant-Laflamme *et al.* (2005) did not ascertain the menstrual cycle phase of their subjects, and hence, were not able to make inferences regarding the effect of sex hormones on the cardiac responses to pain. In addition, a meta-analysis into sex differences in HRV was conducted by Koenig and Thayer (2016), and it was concluded that HRV in females contains more power in the high frequency component (the effect of stress on HRV was not included in the investigation).

The effect of oestrogen on the action of catecholamines has been widely investigated in mammals. In a review of the effect of oestrogen on the stress response, Ueyama *et al.* (2008) reported that HR increases which were induced by stress in ovariectomised rats were greater than those experienced by ovariectomised rats who were supplemented with oestrogen. Ueyama *et al.* (2008) hypothesised that their results were due to oestrogen reducing the sympathoadrenal outflow of stress hormones from the central nervous system. Ueyama *et al.* (2008) also reported that oestrogen reduced the reactivity of the heart to catecholamines, protecting the heart from the effects of stress. If applied to humans, these findings from rats would explain why females in this present study experienced smaller increases in HR when stressed.

Furthermore, the results from the spectral analyses in this study also reveal sex differences. Not only did the males and females exhibit contrasting changes in LF and HF, but the changes in HF were considerably larger than the corresponding changes in LF (see Table 1). The overall stress-induced decrease in LF and the increase in HF in the females is a finding that contradicts the conventional understanding of the relationships between stress and the low frequency and high frequency components of HRV. As already mentioned, studies such as Berntson *et al.* (1994) have previously indicated a lack of consistency in the LF and HF responses to stress after comparing their findings from all-female study cohorts to findings from studies with male cohorts. It is notable that these conclusions were drawn at a time when there was little awareness of sex differences in cardiac dynamics. Therefore, irrespective of the physiological interpretations of LF and HF, the opposing LF and HF trends in males and females, discovered in our pilot study, suggest that the inconsistencies reported by Berntson *et al.* (1994) were probably due to sex differences, and not a redundancy in LF and HF as stress metrics.

It can also not be ignored that the controversy over the physiological interpretation of LF and HF remains largely unresolved. In summary, LF has been speculated to: (i) represent the modulation of both the SNS and PNS (Malik *et al.*, 1996), (ii) be influenced by the frequencies of slow breathing (Brown *et al.*, 1993), and (iii) be influenced by the frequency of muscle contractions in blood vessels (Kenwright *et al.*, 2009).

Similarly, HF has also been suggested to be influenced by typical respiratory frequenc ies (Kenwright *et al.*, 2009). In conclusion, the physiological interpretation of LF and HF cannot be verified without conducting an extensive endocrinological study into the relationship between mental stress, LF, and HF, in which respiration is controlled (Stacey *et al.*, 2018). Also, the spectral results reported here support the LF and HF normalisation method employed in this study. The more common normalisation methods are shown in Eqns. 4 and 5 below.

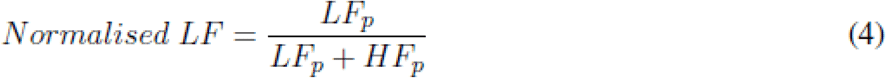

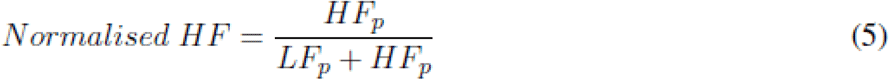

Observe that both of these normalisations would contain the same information, as in this way LFp =1-HF_p_ (Burr, 2007). In contrast, Fig. 1 B and Fig. 1 C establish that the normalised LF and HF metrics proposed in this study contain different information, whereby relatively small changes in LF can be seen alongside large changes in HF. It is evident that the analysis of HRV via LF and HF provides an additional degree of freedom, in comparison to the single degree of freedom offered by HR analysis; without the two degrees of freedom, the sex-specific trends in LF and HF would not have been seen.

The results from the nonlinear analyses in this study also shed new light on the sex differences in the dynamics of ECG. Bornas *et al.* (2006), Vuksanovic and Gal (2007), Williamon *et al.* (2013) and Chanwimalueang *et al.* (2016) have all reported that mental stress causes decreases in the entropy of cardiac signals, however, our results in Fig. 1 D demonstrate that stress caused increases in the entropies computed from the female subjects. These results contradict the complexity loss theory from Goldberger *et al.* (2002), possibly supporting their view that entropies are not true measures of physiological complexity. Nevertheless, the decreased regularity of the HRV from the female subjects confirm that they experienced a stress response which differed from that of the males.

A female-specific stress response has been suggested by Taylor *et al.* (2000), who hypothesised that whilst males experience the conventional fight-or-flight response to stressors, females exhibit a tend-and-befriend response in which they employ social coping methods to combat stress. The tend-and-befriend response is driven by the action of oestrogen and oxytocin (Taylor *et al.*, 2000). Given the effects of oestrogen on cardiac dynamics, the results from this study reveal, for the first time, cardiac trends which are likely to be specific to the tend-and-befriend response. It can be inferred that the fight-or-flight response is characterised by a large increase in HR, an increase in LF, a decrease in HF, and a decrease in HRV entropy; on the other hand, the tend-and-befriend response is characterised by a smaller increase in HR, a decrease in LF, an increase in HF, and an increase in HRV entropy, as illustrated in Figure 2.

**Figure 2.**
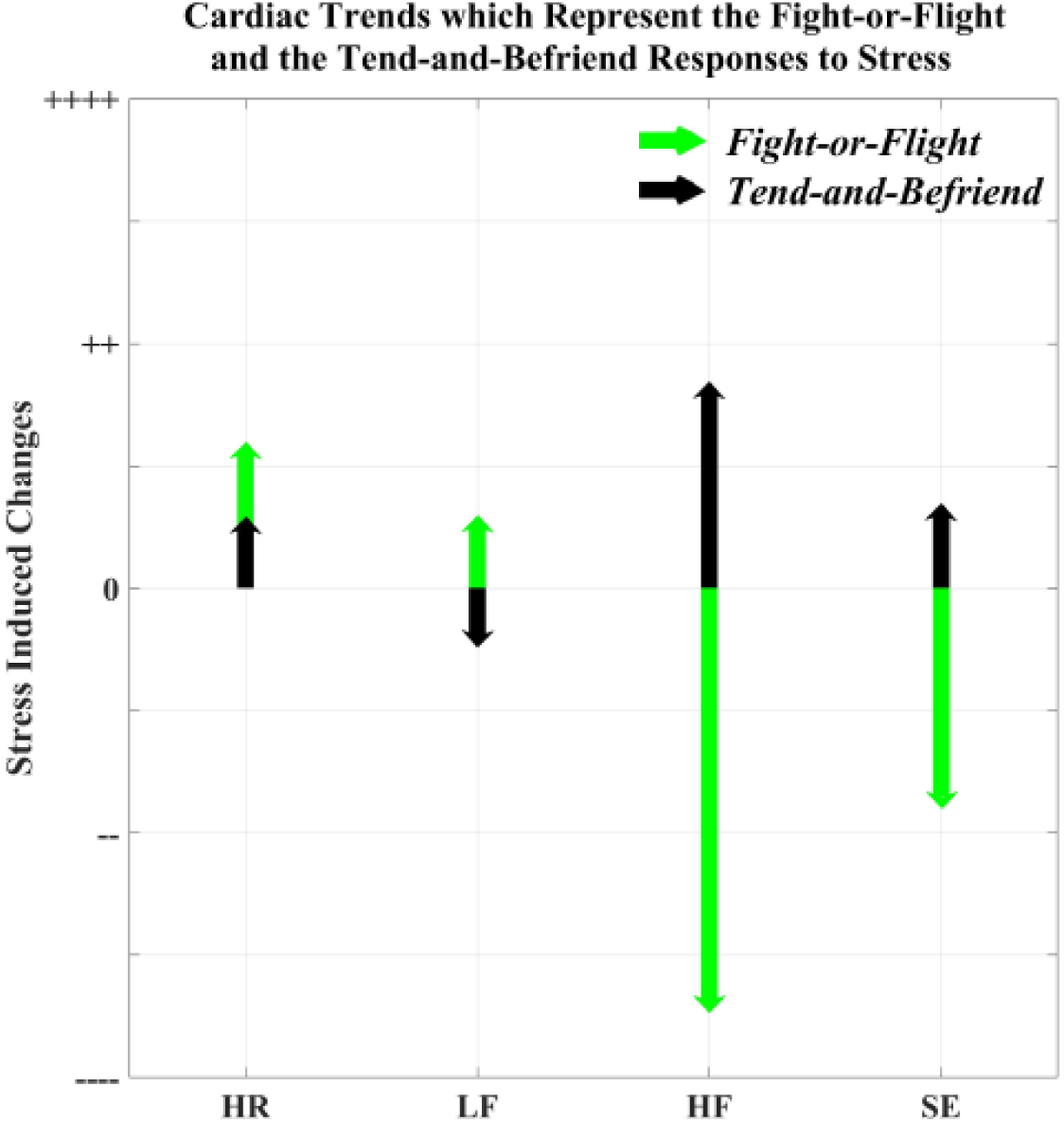
Cardiac trends representing the *fight-or-flight* and the *tend-and-befriend* responses, derived using the results reported in this study.

Future studies will also incorporate endocrinological responses from the subjects, as the use of the calendar method to determine the menstrual phases of the female subjects, while appropriate for this pilot study, does not enable the validation of oestrogen levels. This, in addition to the expansion of the female cohort to include females in the luteal phase, would enable a future comprehensive investigation into the female stress response.

## 5 CONCLUSION

We have investigated long-overlooked sex differences in the cardiac response to stress through temporal, spectral and nonlinear signatures of mental stress in heart rate variability (HRV) signals recorded from 10 females, in the follicular phase of the menstrual cycle, and 10 males.

The cardiac responses were acquired during 15 minutes of rest and 15 minutes of a mental maths test, and the cardiac metrics include the normalised powers of the low frequency component, the normalised powers of the high frequency component, and the sample entropies of the HRV signals, accompanied by statistical comparisons. For rigour, we have also employed a new normalisation procedure for the spectral components of HRV.

Every metric analysed has revealed statistically significant differences (p-value<<0.01) between the cardiac responses in the male and female subjects. In the males only, mental stress has been found to induce large increases in heart rate, increases in the power of the low frequency component of HRV, decreases in the power of the high frequency component of HRV, and decreases in the entropy of HRV. These trends have not been found in the females, suggesting that oestrogen modulates the cardiac response to stress in females. Not only do the results presented here radically challenge the practice of producing scientific hypotheses which have not accounted for sex, but the stress-induced increases in the sample entropy of heart rate variability have never before been reported, thus challenging the common assumption that sample entropy is a reliable measure of signal complexity.

The results from this pilot study have established that the stress-induced cardiac trends which are commonly reported (increases in heart rate and the low frequency component of HRV) are the cardiac manifestations of the fight-or-flight response in males, whereas small increases in heart rate and large increases in the high frequency component of HRV may represent the female tend-and-befriend response. Following this proof-of-concept, further studies will employ females in the luteal phase of their menstrual cycle, and the collection of endocrinological parameters.

### Algorithm 1: Sample Entropy

1. A windowed signal, *x*, of lenth *N* is embedded using an enbedding dimension, *m*, to create (*N* − *m* + 1) segments.Each segment, *X*_*m,*_ is of lenth *m.* In this study, *m* was defined as *m* = 2.
2. A toleranced level of *r* = 0.15 *std* where *std* is the standard deviation in the data window, is defined.
3. The maximun difference, *d*_*max,*_, between the scalar components of two consecutive segments, *X*_*m*_ *(i)* and *X*_*m*_ *(j)*, is computed as

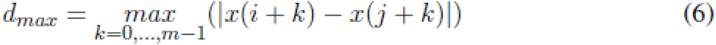
4. For each *X*_*m*_ *(i),* the event *d*_*max*_ < *r* is defined as a match, and a count of such matches is denoted by *A*_*i.*_ The probability of matches, 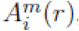, for *X*_*m*_ *(i)* is calculated as

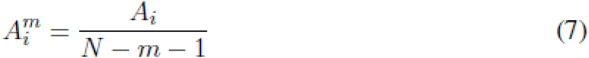 Note: The denominator *N* − *m* − 1, ensure the segment *X*_*m*+1_ (*i*) is included in the computation of the probability.
5. Then the sum of the probabiliy of matches for all segments, Φ, is defined as

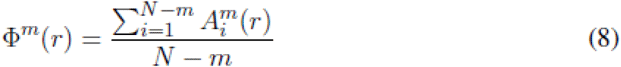
6. The embedding dimenson *m* is increasing to (*m*+1), and step 1 to step 5 are repeated; the sum of the probability of matches for all segment when *m=m*+1 are defined as Ψ and the sample entropy *SE*(*m*,*r*), is computed as

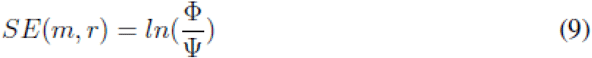
7. The SE for *x* is computed such that one SE value is obtained for each windowed signal.

## AUTHOR CONTRIBUTIONS

The physiological data were recorded by TA and JX, the data analysis was completed by TA, under the supervision of DPM, and the paper was written by TA and DPM.

